# The antifungal activity of cymoxanil is associated with proton pump inhibition and disruption of plasma membrane potential

**DOI:** 10.1101/2024.03.01.583004

**Authors:** Filipa Mendes, Hadar Meyer, Leslie Amaral, Bruno B. Castro, Maya Schuldiner, Maria João Sousa, Susana R. Chaves

**Author notes:** Corresponding author: Susana R. Chaves; E-mail address.

## Abstract

Worldwide use of agrochemicals, particularly pesticides, is necessary to increase agricultural production to feed the ever-growing population. However, despite widespread use, the biochemical mode of action of many agrochemicals and their potential deleterious effects on the environment are poorly characterized. Cymoxanil (CYM) is a fungicide used to combat downy mildew diseases in grapevine cultures and late blight diseases in tomato and potato cultures caused by the oomycetes *Plasmopara viticola* and *Phytophthora infestans*, respectively. Previous reports indicate that CYM affects growth, DNA and RNA synthesis in *Phytophthora* and inhibits cell growth, biomass production and respiration rate in the well-characterized fungal model *Saccharomyces cerevisiae*. We therefore used this model to further dissect mechanisms underlying the toxicological effects of CYM. We found that CYM induced genome-wide alterations, particularly in membrane transporter systems. These alterations were associated with perturbations in lipid-raft organization and inhibition of Pma1p, leading to a decrease in plasma membrane potential and intracellular acidification. Altogether, these findings identify the plasma membrane as one of the targets of CYM and proposes a mode of action underlying its antifungal activity.

## 1. Introduction

Over the last few centuries, the human population has consistently increased and is expected to continue expanding. As a result, agricultural systems have sought increased optimization of productivity, particularly through intensification of the use of agrochemicals (1,2). As pesticides play a pivotal role in the protection of seeds and crops from undesirable organisms, they have been increasingly used as an effective and economical strategy to enhance crop yield and quality (3,4). Pesticides are classified into distinct groups based on the target organism, such as herbicides to control weeds, insecticides to manage insects, rodenticides to control rodents or fungicides to combat fungal diseases (5).

According to the Food and Agriculture Organization of the United Nations (FAO), nearly 4 million tons of pesticides were utilized globally in agriculture in 2021, with fungicides and bactericides accounting for approximately 25 percent of the total^1^. However, the impact of these substances is often overlooked when compared to the effects of insecticides or herbicides (6). While the biochemical modes of action of many fungicides are well characterized, in some cases knowledge is lacking. Indeed, many fungicides are commercially available despite an unknown biochemical mode of action, since fungicide registration often relies solely on fulfilling toxicological and environmental requirements (7). Cymoxanil (CYM) [(1E)-2-(ethylcarbamoylamino)-N-methoxy-2-oxoethanimidoyl cyanide] is one of these.

CYM is a synthetic acetamide compound mainly used in foliar applications in a curative and protective manner (8). It is labelled as a fungicide and is considered crucial to protect grapes and vegetable crops against fungus-like pathogens, namely oomycete pathogens from the Peronosporales order (9), downy mildew diseases caused by *Plasmopara viticola* in grapevine cultures, and late blight diseases triggered by *Phytophthora infestans* in tomato and potato crops (10,11). Given its relatively limited period of efficiency, CYM is frequently paired with other fungicides such as mancozeb and copper, known for their multisite mode of action and extended protective effects, thus increasing overall efficacy (12). However, only a few studies have addressed its mode of action. It is known that CYM inhibits mycelial growth and germ tube formation by sporangia in *P. infestans*. CYM also inhibits DNA and RNA synthesis, particularly the former, but these alterations are considered secondary effects and not its primary mode of action (13). CYM can also affect non-target organisms, as it can inhibit growth of the bacteria *Rhodopirellula rubra* and the microalgae *Raphidocelis subcapitata* (8). Even though the toxicological profile of CYM has been poorly characterized, the European Commission still renewed the authorization to use CYM and products containing CYM until 2026. It is therefore essential to reinforce studies to uncover its mode of action.

The yeast *S. cerevisiae* has previously been used to further dissect potential cellular effects of CYM, which resulted in inhibition of cell growth, biomass production and respiration in one yeast strain (14) and reduced ATP synthesis in another (9). Although it is not a CYM target organism, this well-characterized model system presents several advantages. On one hand, it shares essential cellular pathways affected by CYM with higher eukaryotes, humans included, facilitating the understanding of the fungicide’s impact in both target and non-target organisms. Moreover, its genetic tractability, functional information available for nearly every gene, and availability of a vast array of dedicated experimental tools, protocols, software and databases, facilitates the identification of specific targets and pathways affected by CYM, providing insights into its mode of action (15,16). Additionally, the efficiency and cost-effectiveness of working with *S. cerevisiae* streamlines experimental procedures, allows high-throughput studies integrating genome-wide methodologies and conventional toxicological investigations. Indeed, omics-based approaches provide a comprehensive evaluation and characterization of molecular mechanisms and the cellular response to several toxic substances at the genome, transcriptome, proteome, and metabolome levels (17). In this context, *S. cerevisiae* has been used to identify mechanisms of action and potential targets of new drugs (18–20) and has proven invaluable as a first screening tool, limiting the use of animal models. Taking all of the above into consideration, we decided to explore this well-characterized model system to further dissect mechanisms underlying the toxicological effects of CYM.

## 2. Materials and methods

### 2.1. Yeast Strains

The yeast *S. cerevisiae* BY4741 (MAT a *his3Δ1; leu2Δ0; met15Δ0; ura3Δ0*) was used throughout this study as the wild-type strain. The *end3Δ* mutant in the same background was obtained from the Euroscarf collection. BY4741 cells were transformed with the pRS416 GFP-Atg8 plasmid (21) using the LiAc/SS Carrier DNA/PEG method (22).

### 2.2. Growth Conditions and treatments

BY4741 and *end3Δ* were cultured in Synthetic Complete medium [SC; 0.5 % (w/v) ammonium sulfate; 0.17% (w/v) yeast nitrogen base without amino acids and without ammonium sulphate; 0.14% (w/v) drop-out mixture lacking leucine, histidine, tryptophan, and uracil; 0.04% (w/v) leucine; 0.008% (w/v) tryptophan; 0.008% (w/v) histidine, and 0.008% (w/v) uracil] with 2% of glucose (Glu) or galactose (Gal) as the carbon source. The BY4741 strain expressing pRS416 GFP-Atg8 was grown in SC medium with glucose, but lacking uracil. All yeast strains were grown at 30 °C in an orbital shaker at 200 rpm, with a ratio of flask volume/medium of 5:1.

For all assays, cells were grown overnight, then diluted in fresh medium to OD_640 nm_ = 0.1 and treated with CYM (25 or 50 μg mL^-1^, as indicated in the figure legends) from Sigma-Aldrich and/or the equivalent volume of DMSO as a negative control (lower than 0.5 %) for up 8 h. Samples were taken at specific time points for subsequent analyses. Additionally, samples of wild-type strain BY4741, grown in glucose and treated for 8 h, were collected for transcriptomic analysis, evaluation of lipid raft distribution, quantitative Real-Time PCR (qRT-PCR), analysis of efflux pumps, and assessment of proton movements, intracellular ATP concentration, vacuolar pH variations and plasma membrane potential. The evaluation of lipid raft distribution was also assessed at the *end3Δ* mutant. BY4741 strain harboring pRS416 GFP-Atg8 was used to assess autophagy.

### 2.3. Cell growth and survival

Cell growth was assessed by measuring the OD_640nm_ of yeast cells at time point 0, 4 and 8 h in the absence or presence of CYM. Cell survival was evaluated by counting of colony forming units (c.f.u.). Briefly, five serial dilutions of the cultures were performed, and 40 μL of 10^-4^ or 10^-5^ dilutions (according to each condition) were plated onto yeast extract-peptone-dextrose plates [YEPD; 1% (w/v) yeast extract; 2% (w/v) peptone; 2% (w/v) glucose and 2% (w/v) agar] and incubated for 48 h at 30 ºC. The percentage of cell survival was calculated from the number of c.f.u. of each condition in relation to time zero and the control without any treatment.

### 2.4. Assessment of plasma membrane integrity

To assess loss of plasma membrane integrity, cells were stained with the cell-impermeant dye propidium iodide (PI, Sigma-Aldrich). At each timepoint, 500 μl of cells at OD_640nm_ = 0.1 were harvested and resuspended in phosphate buffered saline (PBS) 1× (PBS 10×; 137 mM NaCl, 2.7 mM KCl, 10 mM Na2HPO4, 1.8 mM KH2PO4, pH 7). PI was added to the yeast cell suspension to a final concentration of 2 μg mL^-1^ and incubated for 10 min at room temperature. Cells with red fluorescence (PI positive cells) were considered to have lost their plasma membrane integrity.

### 2.5. Transcriptomic analysis

#### 2.5.1. RNA extraction

Total RNA was extracted from cells using TRIzol/chloroform and RNA Clean & Concentrator-5 Kit (Zymo Research) according to manufacturer’s instructions. Briefly, the upper aqueous phase from TRIzol/chloroform extraction was transferred to a new tube, and RNA was purified with the RNA Clean & Concentrator-5 Kit. The extracted RNA was used for RNA-sequencing (RNAseq) analysis and for qRT-PCR.

#### 2.5.2. RNAseq analysis

RNAseq analysis was performed by the Genomics Facility at the Gulbenkian Institute of Science, as a paid service. Samples were processed with the RNA Lexogen QuantSeq 3’ mRNA-Seq V2 Library Prep Kit FWD with UDI 12 nt. The differential expression gene analysis between CYM-treated cells and non-treated cells was performed with the package DESeq2. The threshold for up-regulated genes as log2 fold change > 2 and adjusted P value < 0.05 and for down-regulated genes as log2 fold change < -2 and adjusted P value < 0.05, the genes satisfying these criteria were identified as differentially expressed genes.

#### 2.5.3. qRT-PCR

cDNA was synthesized by reverse transcription from 500 ng of total RNA according to manufacturer’s instructions (iScript™ cDNA Synthesis Kit, Bio-Rad Laboratories). Real-time PCR reactions using primers hybridizing with *ERG25* or *ACT1* transcripts were performed in an CFX96^TM^ Real-Time System C100^TM^ Thermal Cycler (Bio-Rad Laboratories) using the KAPA SYBR® FAST qPCR Master Mix (2×) Kit (Sigma-Aldrich) according to manufacturer’s instructions. PCR controls with no template were also performed for each primer pair. Two biological replicates were performed, and each sample was measured in duplicate. Quantification was performed using the 2^-ΔΔCt^ method (23) and expression of *ERG25* was normalized to A*CT1*.

### 2.6. High-throughput fluorescence microscopic screen

A high-throughput fluorescence microscopy screen was performed using a collection of 5,330 strains containing an N-terminal SWAT cassette harboring the *NOP1* constitutive promotor and GFP upstream of each gene start codon (N′ NOP1pr-GFP library) (24,25). The strains were grown in Synthetic Defined medium [SD; 0.67% (w/v) yeast nitrogen base without amino acids [Conda Pronadisa] and 2% (w/v) dextrose, lacking uracil]. Cells were moved from agar plates into liquid 384-well polystyrene growth plates using the ROTOR HDA (Singer Instruments). Liquid cultures were grown overnight in 50 μl of SD medium in a shaking incubator (LiCONiC Instruments) at 30 °C. Afterwards, 5 μl of overnight culture was back-diluted into 95 μl of SD medium with 25 μg mL^-1^ of CYM and/or the equivalent volume of DMSO as a negative control (lower than 0.5%), using a Tecan freedom EVO liquid handler (Tecan) connected to the incubator. Plates were then transferred back to the incubator and grown for 8 h at 30 °C. The liquid handler was then used to transfer strains into glass-bottom 384-well microscope plates (Brooks Bioscience) coated with Concanavalin A (Sigma-Aldrich) to allow cell adhesion. Wells were washed twice in SD medium to remove floating cells and obtain a cell monolayer. Plates were then transferred into an automated microscopy system using a KiNEDx robotic arm (Peak Robotics).

Imaging was performed with an automated Olympus SpinSR system using a Hamamatsu flash Orca 4.0 camera and a CSUW1-T2SSR SD Yokogawa spinning disk unit with a 50 μm pinhole disk. Images were acquired using a 60× air lens NA 0.9 (Olympus), laser excitation 488nm, intensity 50% and exposure time 500 ms, with CSU dichroic mirror D405/488/561 and emission filter set B525/50. Images were manually inspected using ImageJ software.

### 2.7. Gene ontology (GO) term analysis

GO term enrichment analysis was performed using the *Saccharomyces* Genome Database (SGD) GO Term Finder. The background set for each analysis (RNAseq and microscopy) was defined as all the genes for which data was available. A *p* value < 0.05 was considered statistically significant.

### 2.8. Evaluation of autophagy

Autophagy was assessed by western blot analysis of GFP-Atg8 cleavage. Atg8p is incorporated into the autophagosome and later degraded in the vacuolar lumen (26); however, GFP is resistant to degradation in the vacuole and thus free GFP detected by western blot in cells expressing GFP-Atg8 is indicative of autophagy (21). At each timepoint, 1 mL of BY4741 cells harboring pRS416 GFP-Atg8 cells at an OD6_40nm_ = 1 were harvested by centrifugation at 5000 ×g for 3 min and washed once with distilled water. For preparation of whole cell extracts, cells were then resuspended in 500 μL of water containing 50 μL lysis buffer (3.5% (v/v) β-mercaptoethanol in 2 M NaOH) and incubated for 15 min on ice. Next, 50 μL of 3 M trichloroacetic acid were added and tubes incubated for 40 min on ice to precipitate the proteins. Extracts were centrifuged at 12000 ×g for 10 min at 4 °C, washed with 100 μL of acetone and centrifuged again at 12000 ×g for 5 min at 4 °C. Finally, protein extracts were resuspended in Laemmli buffer (2% (v/v) β-mercaptoethanol, 0.1 M Tris pH 8.8, 20% (v/v) glycerol, 0.02% (v/v) bromophenol blue). Samples were heated at 70 °C for 15 min and stored at -20 °C, or were used immediately for western blot.

Samples were separated electrophoretically on a 12.5% sodium dodecyl sulphate-polyacrylamide gel (SDS-PAGE) and transferred to Polyvinylidene Difluoride (PVDF, GE Healthcare) membranes at 60 mA per membrane for 90 min in a semi-dry transfer unit (TE77X Hoefer). To avoid non-specific interactions, membranes were blocked in 5% non-fat milk in PBS-Tween 0.1% (v/v) solution (1× PBST) with agitation at RT for 1 h. Afterwards, membranes were incubated overnight at 4 °C with anti-GFP (1:3000, Sigma-Aldrich) and anti-yeast phosphoglycerate kinase (Pgk1p) mouse monoclonal primary antibodies (1:5000, Molecular Probes). Pgk1p was used as the loading control. The following day, membranes were incubated with the Peroxidase-AffiniPure goat anti-mouse IgG secondary antibody (1:5000, Jackson ImmunoResearch). Chemiluminescence detection was performed using an ECL detection system (Millipore-Merck) in a G:BOX Chemi XX9 system (Syngene).

### 2.9. Evaluation of lipid raft distribution

The distribution of lipid rafts was inferred by fluorescence microscopy (Leica Microsystems DM-5000B epifluorescence microscope) using filipin staining (Filipin III from *Streptomyces filipinensis*, Sigma-Aldrich) as previously described (27,28). Filipin is a naturally fluorescent antibiotic dye that binds to ergosterol and thus is used to detect regions with high sterol content in the plasma membrane of fungal species (29). Briefly, cells were collected at an OD_640 nm_ = 0.5 and concentrated 20× in sterile water. Immediately before visualization, cells were incubated for 1 min in the dark with 0.1 g L^-1^ filipin from a stock solution of 5 g L^-1^ (w/v) in methanol. Cells were then mounted on slides with the anti-fading agent Vectashield (Vector Laboratories) to overcome the instability of this dye, and then visualized in an epifluorescence microscope.

### 2.10. Analysis of efflux pumps

To assess presence of efflux pumps cells were stained with the cell-permeant dye calcein AM (Molecular probes), as previously described (30). Briefly, cells were harvested (500 μl at OD6_40nm_ = 0.1), resuspended in 1× PBS and incubated with 100 μM of verapamil, a known inhibitor of fungal efflux pumps (31–33), for 90 min at room temperature. Then, 10 μM of calcein AM, a cell-permeant and non-fluorescent dye that is cleaved by intracellular esterases into green-fluorescent calcein, which is retained in the cytoplasm (34,35), were added to both verapamil-treated and untreated cells. After 30 min of incubation at room temperature, cells were analyzed by flow cytometry. Data are presented as the ratio of between the median green fluorescence intensity of verapamil-treated and untreated cells (+ verapamil/ - verapamil).

### 2.11. Proton movements assays

Cells were harvested (3 mL of cells at OD_640nm_ = 20), washed and resuspended in distilled water. Proton movements were measured at room temperature by recording the pH of cell suspensions with a standard pH meter (PHM 82; Radiometer) connected to a potentiometer recorder (BBC-GOERZ METRAWATT, SE460). The pH electrode was immersed in 4.5 mL water and 0.5 mL of yeast suspensions with magnetic stirring. For measurement of proton efflux, the initial pH was adjusted to 5.5 and a baseline was established. Afterwards, 20 mM of glucose was added to trigger H+ efflux, leading to an acidification of the extracellular environment (36). The rate of acidification, calibrated with 0.1 mM NaOH, was taken as a measure of proton extrusion activity. A sample was taken from each condition to estimate the dry weight (DW) after incubation at 80 °C for two days. Data is presented as Protons min^-1^ mg^-1^ DW.

### 2.12. Evaluation of intracellular ATP concentration

Intracellular ATP was quantified using the BacTiter-Glo™ Microbial Cell Viability kit as recommended. Cells were transferred to a 96-well opaque plate, mixed with an equal volume of BacTiter-Glo reagent and incubated for 15 min in the dark. The emitted luminescence was recorded in a Varioskan flash multimode reader (1000 ms integration time). Data are shown in arbitrary units as fold change expressed in relation to untreated cells (control, considered as 1).

### 2.13. Analysis of vacuolar pH variations

Vacuolar pH alterations were detected using the 5-(and-6)-carboxy-2′,7′-dichlorofluorescein diacetate (CDCFDA, Invitrogen) probe. CDCFDA, a non-fluorescent cell permeant dye, is cleaved by intracellular esterases and yields the fluorescent and pH-sensitive 5-(and-6)-carboxy-2’,7’-dichlorofluorescein (CDCF). At higher pH values, CDCF accumulates in the yeast vacuole, and cells exhibit higher fluorescence intensity (37,38). Cells were harvested by centrifugation (0.5 mL OD_640nm_ = 0.1), washed once with sterile water and resuspended in CF buffer [50 mM glycine, 10 mM NaCl, 5 mM KCl, 1 mM MgCl2, 40 mMTris, 100 mM MES pH 4.5 supplemented with 2% (w/v) glucose] (28,37). The cell suspension was then incubated with 1.6 μM CDCFDA for 20 min at 30 °C with 200 rpm orbital agitation. Afterwards, cells were washed and resuspended in CF buffer without glucose for further analysis in the flow cytometer. Flow cytometry data are presented as the median of the green fluorescence intensity in relation to the autofluorescence of each sample and untreated cells.

### 2.14. Assessment of membrane potential

Plasma membrane potential was monitored using Bis-(1,3-dibutylbarbituric acid) trimethine oxonol [DiBAC_4_(3), Molecular Probes]. Normally, this dye is excluded by polarized cells, but accumulates inside depolarized cells. Therefore, increased fluorescence is indicative of loss of plasma membrane potential (39). Cells were harvested by centrifugation (0.5 mL OD_640nm_ = 0.1) and resuspended in 1× PBS. DiBAC_4_(3) and PI were added to the yeast cell suspension to a final concentration of 1 μg mL^-1^ and 2 μg mL^-1^, respectively, and samples incubated for 10 min at room temperature and analyzed by flow cytometry. Data are presented as percentage of cells with green fluorescence [DiBAC_4_(3) positive cells] and without red fluorescence (PI negative cells), corresponding to depolarized but non-permeabilized cells.

### 2.15. Flow cytometry

Flow cytometry analysis was performed with CytoFLEX (Beckman Coulter Inc.) with 488 nm emission. Filter set included a 525/40 nm bandpass filter for ‘‘green’’ fluorescence FITC channel and a 585/42 nm bandpass filter for ‘‘red’’ fluorescence PE channel. For each sample, 10,000 events were evaluated. Data were analyzed using CytExpert software.

### 2.16. Statistical analysis

The statistical analysis of the results was performed using GraphPad Prism software. In all assays, data are expressed as the mean and standard deviation (S.D.) and a one-way analysis of variance (ANOVA) was used to test the effect of CYM and a Dunnett test was used to assess differences relatively to control. A significance level of 0.05 was employed in all analyses.

## 3. Results

### 3.1. Cymoxanil inhibits growth and decreases cell viability independently of an active respiratory chain

To ascertain if the cytotoxic effect of CYM in *S. cerevisiae* is due to its previously described effect in inhibiting respiration, we directly compared how it affects growth and viability of cells in medium containing glucose or galactose as the carbon source, where respiration is repressed or not, respectively. As seen in Figure 1A, CYM inhibited growth of yeast cells independently of the carbon source, but a significant viability decrease was observed after 4 h of treatment in glucose (Figure 1B, blue bars), but only after 8 h of treatment in medium containing galactose (Figure 1B, orange bars). Cells exposed to CYM in glucose-containing medium also lost plasma membrane integrity earlier, as staining with PI was already visible after 4 h, and the percentage of cells with a compromised plasma membrane after 8 h was much higher in glucose-than galactose-grown cells (Figure 1C). Taken together, our data indicates that cells are more sensitive to CYM in medium containing glucose, and we therefore pursued further characterization of the effects of CYM in this medium.

**Figure 1.**
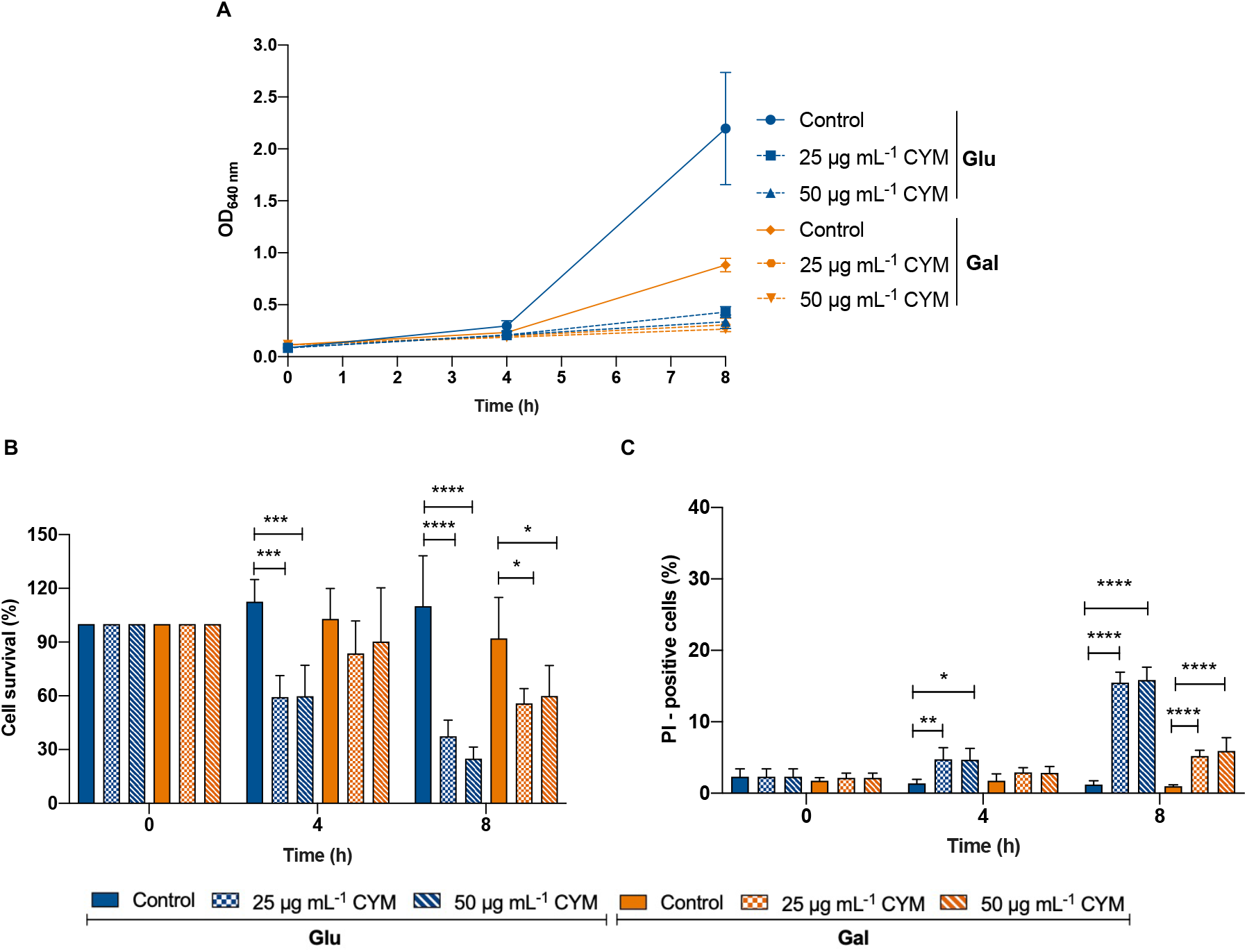
Influence of the carbon source on CYM effects on growth and viability of *S. cerevisiae* BY4741 cells. Cells were grown overnight in SC medium containing 2% glucose (Glu) or 2% galactose (Gal) then were diluted to an OD_640nm_ = 0.1, transferred to fresh medium and treated with 25 or 50 μg mL^-1^ CYM or the equivalent volume of DMSO (control). **(A)** OD_640nm_ were registered at time point 0, 4 and 8 h of treatment. **(C)** Cell survival of CYM-treated cells was determined by standard dilution plate counts and expressed as a percentage of c.f.u. on YEPD plates in relation to time 0. **(D)** Plasma membrane integrity was determined by PI staining of cells treated with 0, 25 and 50 μg mL^-1^ CYM at time point 0, 4, 8 h of treatment in SC medium containing glucose or galactose. The data displayed are the mean ± standard deviation of three independent experiments. Asterisks (* ≤ 0.05; **, P ≤ 0.01; ***, P ≤ 0.001; ****, P ≤ 0.0001) depict significant differences relative to the control at each time point.

### 3.2. Transcriptome alterations induced by cymoxanil

To uncover the global cellular effects of CYM on yeast cells, we used two non-biased approaches: RNAseq to determine how this fungicide affects the RNA level of yeast genes and a high-throughput microscopic screen to assess its effects on protein localization.

Transcriptomic analysis performed in cells incubated in the presence of 25 μg mL^-1^ CYM, for 8 h, indicated that a total of 70 genes were differentially expressed when compared with cells incubated with the CYM vehicle, DMSO, for the same time. From these, 50 were up-regulated, whereas 20 were down-regulated, considering a threshold of log2 fold change > 2 and adjusted *p* value < 0.05 and log2 fold change < -2 and adjusted *p* value < 0.05, respectively (Figure 2A). The top 10 up-regulated genes and top 10 down-regulated genes are shown in Figure 2B. GO enrichment showed that sulfur metabolism, oxoacid metabolism and organic acid metabolism were the biological functions up-regulated by CYM (Figure 2C), whereas one-carbon, glycine, small molecule, and serine family amino acid metabolism were those most enriched in the dataset of down-regulated genes (Figure 2D). In both data sets, cell periphery was the most enriched cellular component (Figure 2C and 2D).

**Figure 2.**
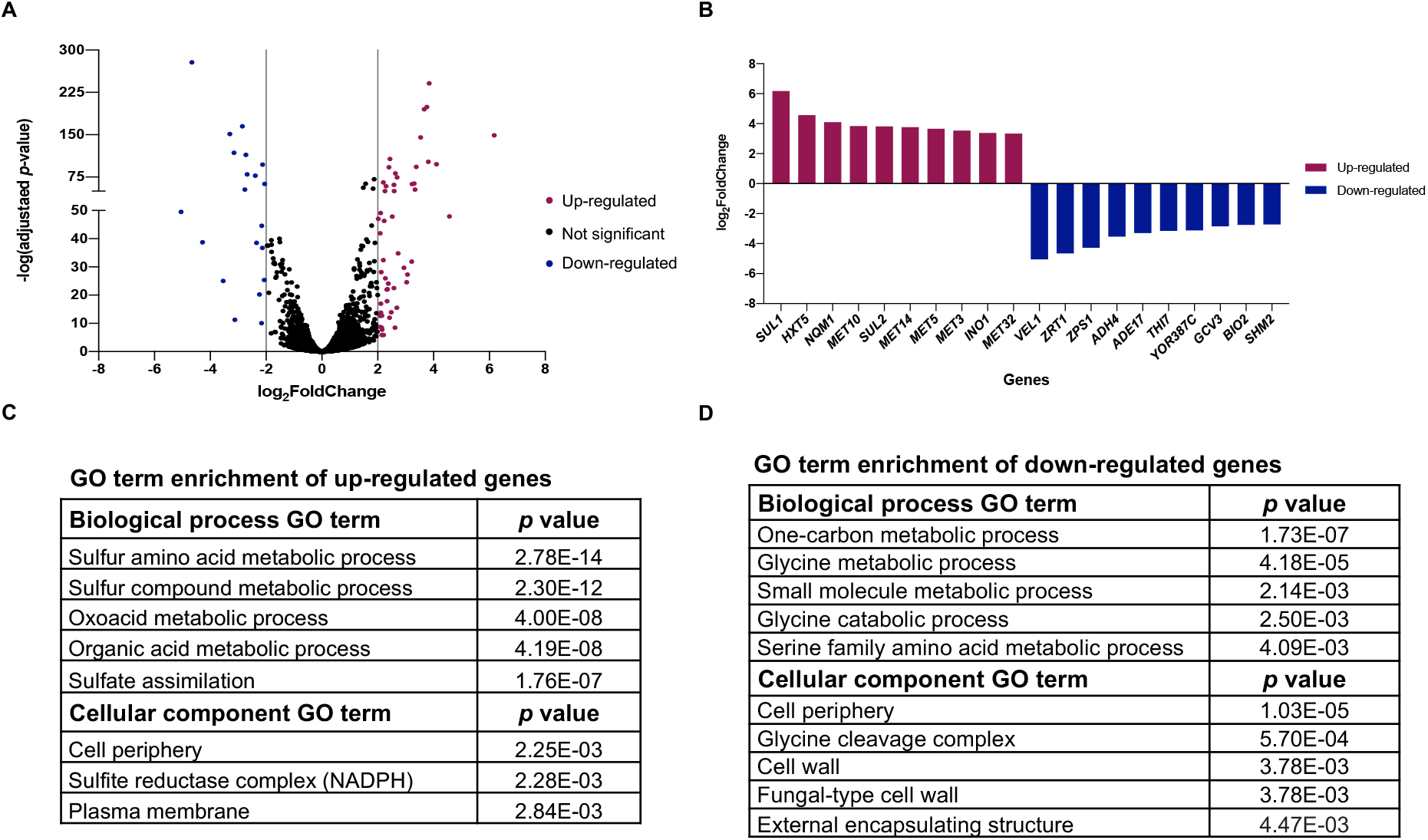
Transcriptomic analysis of CYM effect in *S. cerevisiae* BY4741 cells. Cells were treated for 8 h in the absence (control) or presence of 25 μg mL^-1^ of CYM, and RNA was purified for RNAseq analysis. (**A**) Volcano plot of the differentially expressed genes. The x-axis represents log2Fold change and the y-axis represents -log10(adjusted *p*-value). The purple points represent up-regulated genes, and blue points represent down-regulated genes. Genes not significantly altered are shown in black. The threshold for differential expression was set as log2 fold change > 2 and log2 fold change < -2, and adjusted *p* value < 0.05. (**B**) The top 10 up-regulated genes (purple) and the top 10 down-regulated genes (blue). The top 5 of GO term enrichment analysis of the 50 up-regulated genes **(C)** and the 20 down-regulated genes **(D)**.

### 3.3. Proteome-wide alterations induced by cymoxanil

To look at possible effects of CYM on the proteome, we imaged a collection of yeast strains each expressing one yeast protein tagged with a GFP at its N terminus under control of a constitutive *NOP1* promoter. The advantage of this collection is that the control of a constitutive promoter allows us to focus on post transcriptional events.

Proteome alterations were assessed in cells incubated in the presence of 25 μg mL^-1^ CYM, for 8 h, and proteins were considered to display altered localization (partially or totally) when compared with cells incubated with the CYM vehicle, DMSO, for the same time (examples in Figure 3A). We found that, of a total of 1648 proteins analyzed, 224 changed their localization.

**Figure 3.**
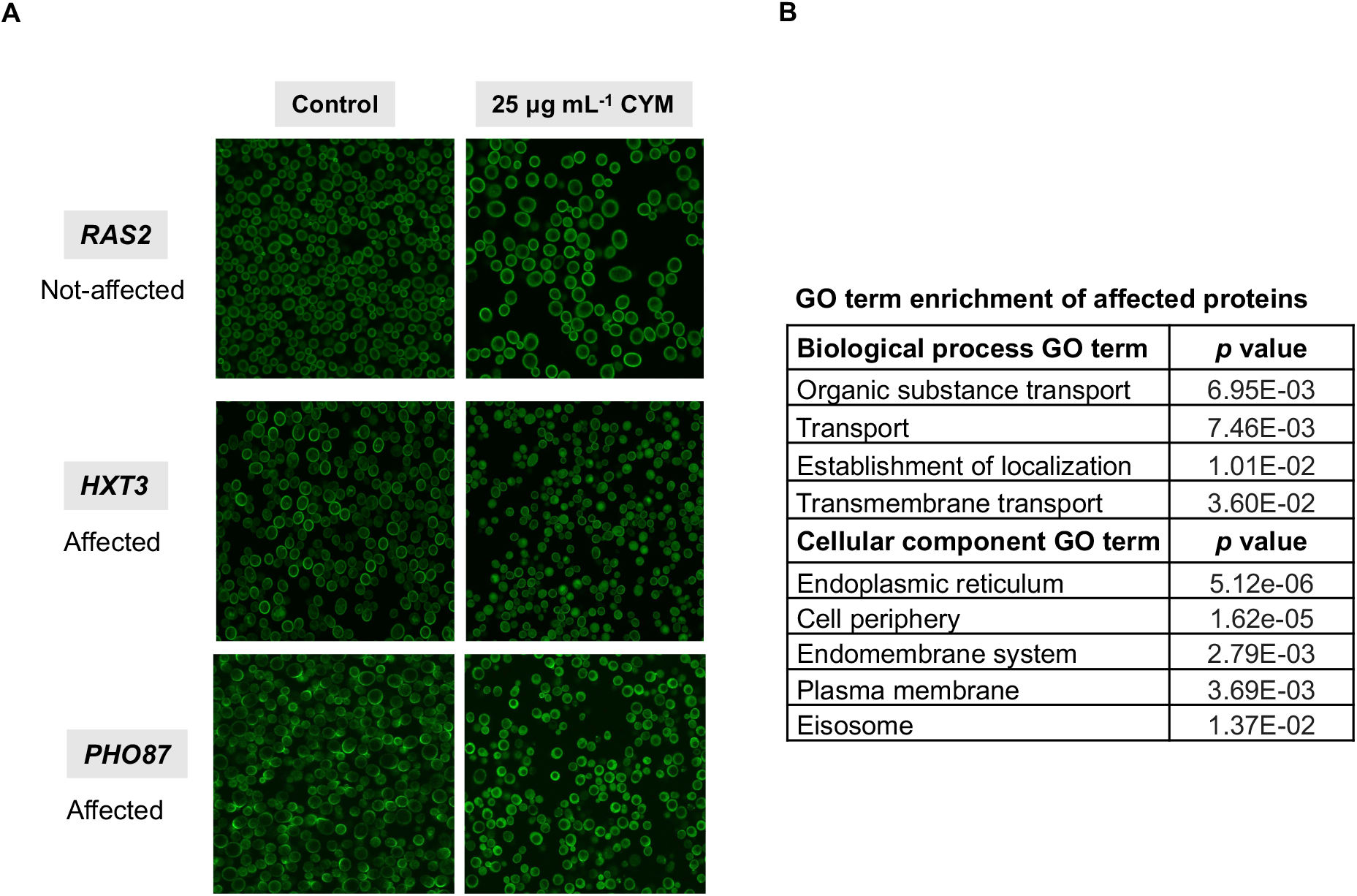
Effects of CYM on protein localization. Yeast strains in the collection (N′ *NOP1*pr-GFP library) were grown overnight in SD liquid medium, back-diluted 1:20 into 95 μl of fresh medium with 25 μg mL^-1^ of CYM and/or the equivalent volume of DMSO as a negative control, and grown for 8 h at 30 °C before image acquisition and manual assessment of the localization of the GFP tagged protein. Proteins for which data was obtained were divided into two groups: proteins that do not change their localization and proteins demonstrating altered localization. **(A)** Representative images from the two groups **(B)** The top 5 of GO term enrichment analysis of proteins affected by CYM.

GO enrichment analysis showed that proteins involved in biological processes related to transport and transmembrane transport were overrepresented in the dataset of proteins whose localization was partially or completely altered by exposure to CYM (Figure 3C). Furthermore, the cellular components with higher representation were the endoplasmic reticulum, with several proteins displaying additional diffuse localization, and cell periphery, with many proteins also accumulating in the vacuole (Figure 3C).

### 3.4. Cymoxanil does not induce autophagy in yeast cells

Results from the two screens seemed to suggest that CYM may induce a response similar to starvation, with increased catabolism and accumulation of several membrane proteins in the vacuole. We thus investigated if these observations could be a result of autophagy induction. However, as seen in Figure 4, CYM did not lead to an increase in GFP-Atg8 delivery to the vacuole and consequent cleavage to free GFP, one of the most commonly used markers to ascertain autophagy in yeast.

**Figure 4.**
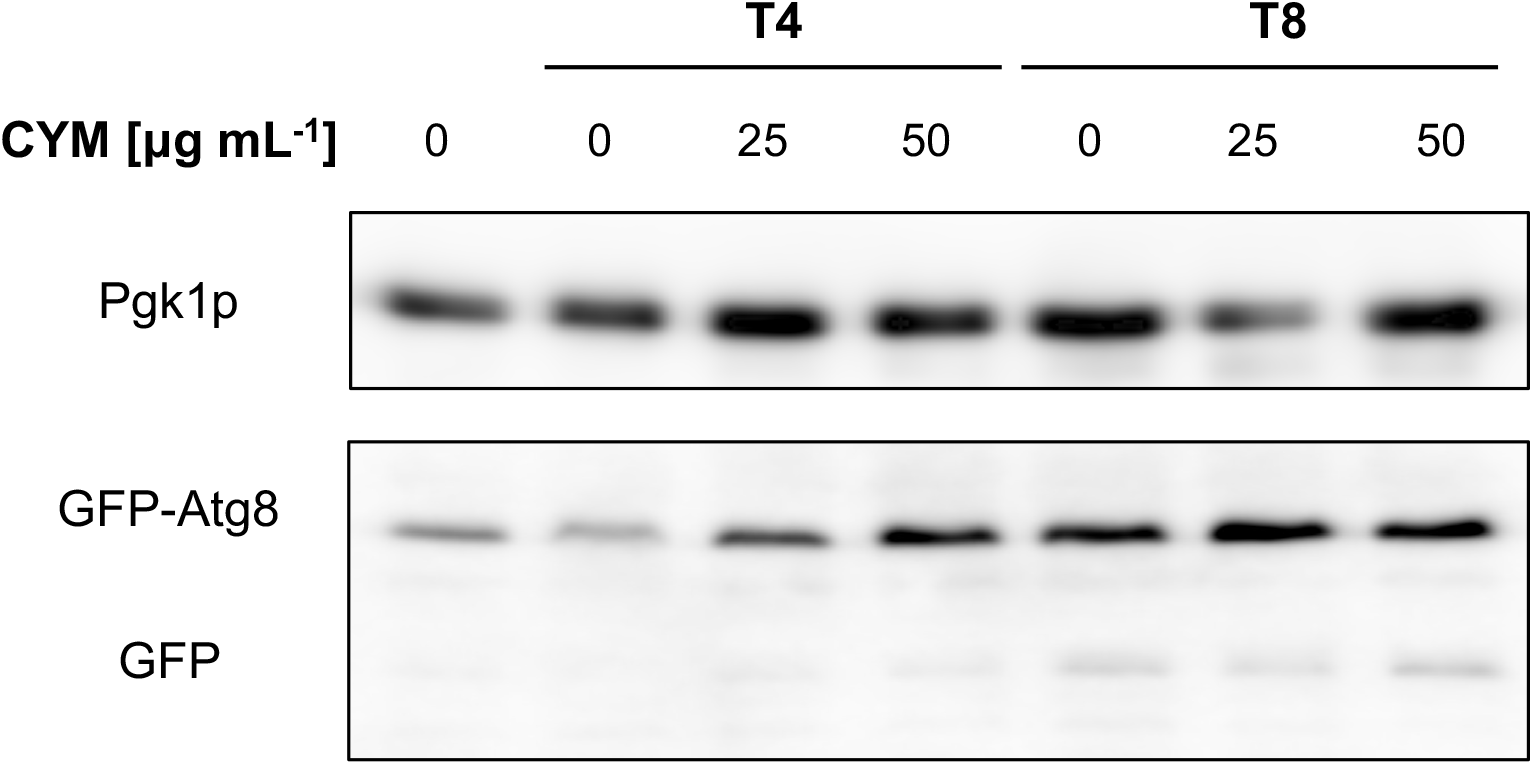
Effect of CYM on autophagy in BY4741 cells. BY4741 cells harboring pRS416 GFP-ATG8 were treated for 8 h in the absence (control) or presence of 25 or 50 μg mL^-1^ of CYM. Samples were collected before (time 0) and after 4 and 8 h of treatment. Autophagy was monitored by western-blot analysis of GFP-Atg8 cleavage. Pgk1p was used as the loading control.

### 3.5. Cymoxanil induces intracellular accumulation of ergosterol, perturbing lipid rafts of yeast cells without affecting ergosterol biosynthesis

Results from both screens also indicated that CYM affects transporter proteins, which may be a direct effect or a result of alterations in the plasma membrane. We therefore assessed if CYM affects the distribution of lipid rafts, domains enriched in ergosterol and sphingolipids, with the fluorescent sterol marker filipin. In untreated cells, a normal punctate pattern of filipin staining in the plasma membrane was observed (arrows), while intracellular spots (arrowheads) were evident in CYM-treated cells (Figure 5A). This phenotype was also observed in the *end3*Δ mutant, which is defective in the internalization step of endocytosis (Figure 5A). These CYM-induced alterations were not a result of ergosterol depletion, which would result in up-regulation of *ERG* genes, which we did not observe by transcriptomic analysis of CYM-treated cells. We further confirmed there was no up-regulation of *ERG25* by qRT-PCR (Figure 5B).

**Figure 5.**
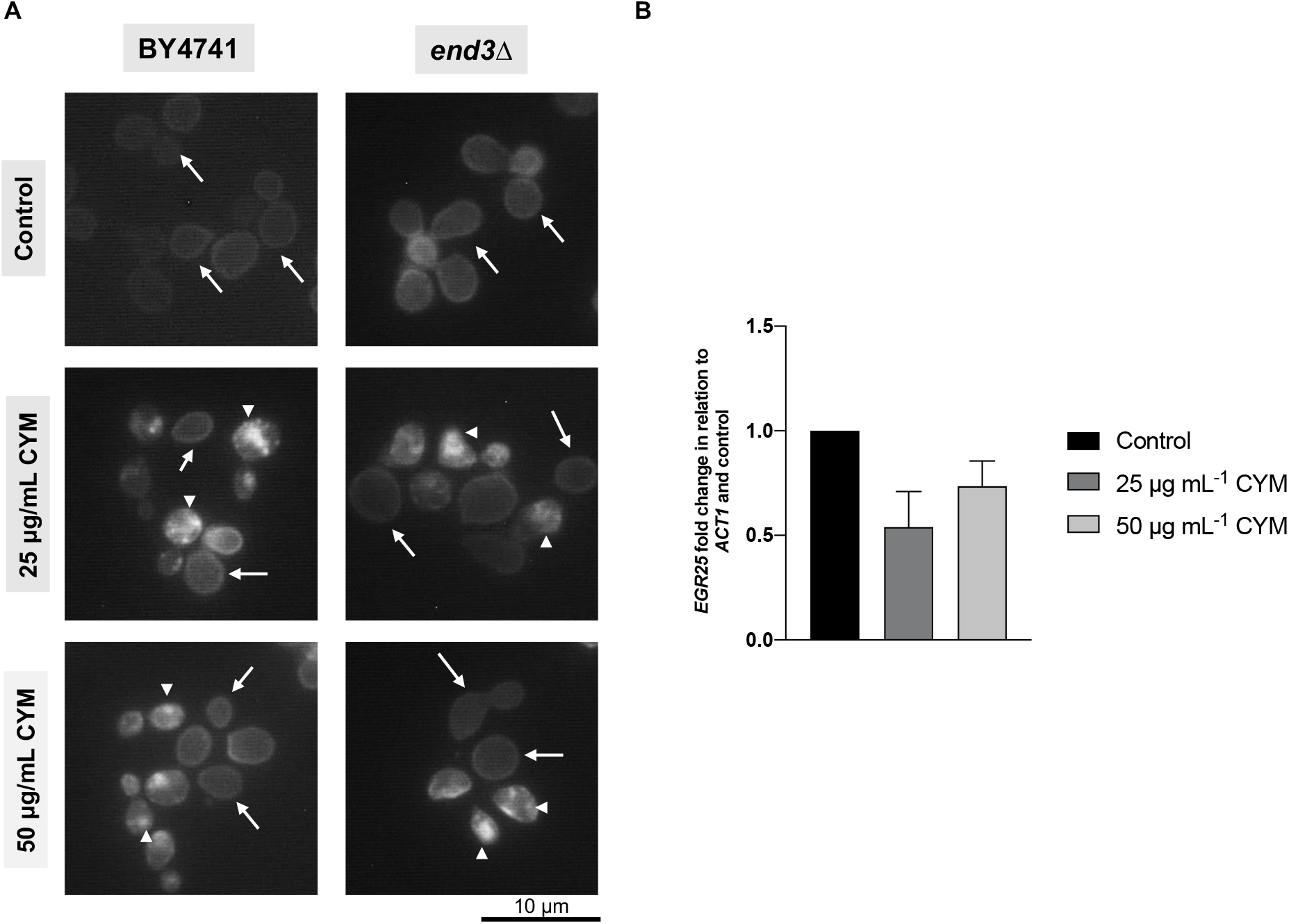
Effect of CYM on the distribution of ergosterol-rich lipid rafts. **(A)** Wild type and *end3*Δ cells were treated for 8 h in the absence (control) or presence of 25 or 50 μg mL^-1^ of CYM. Then, cells were collected and stained in the dark with 0.1 μg L^-1^ filipin, immediately before visualization under the microscope. Representative fluorescence images of each condition are shown. Due to fast photobleaching of filipin, fluorescence intensity is not associated with ergosterol levels and only localization can be inferred. Head arrows: intracellular filipin staining. Arrows: plasma membrane filipin staining. **(B)** Wild type cells were treated 8 h in absence (control) or presence of 25 or 50 μg mL^-1^ CYM and *ERG25* expression levels were analysed by qRT-PCR using *ACT1* as normalizer. The expression values are represented as the fold change relative to the nontreated cells (control) and correspond to the mean ± standard deviation of two independent experiments.

### 3.6. Cymoxanil does not affect efflux pumps but inhibits Pma1p activity and leads to plasma membrane depolarization of yeast cells

As alterations in lipid rafts can influence the activity of proteins that are normally localized in these domains, we next assessed whether CYM can affect efflux pumps. For that purpose, cells were exposed to CYM and then stained with calcein AM in the absence or presence of the efflux pump inhibitor verapamil. If CYM caused an inhibition or decrease of efflux pumps in the membrane, it would be expected that the ratio between the fluorescence intensity of cells that were treated with verapamil or left untreated would be different in CYM-treated cells in comparison with the control. However, this ratio of fluorescence intensity (+ verapamil/ - verapamil) was similar in CYM-treated cells and non-treated cells, indicating that CYM does not affect efflux pumps in yeast cells (Figure 6A).

**Figure 6.**
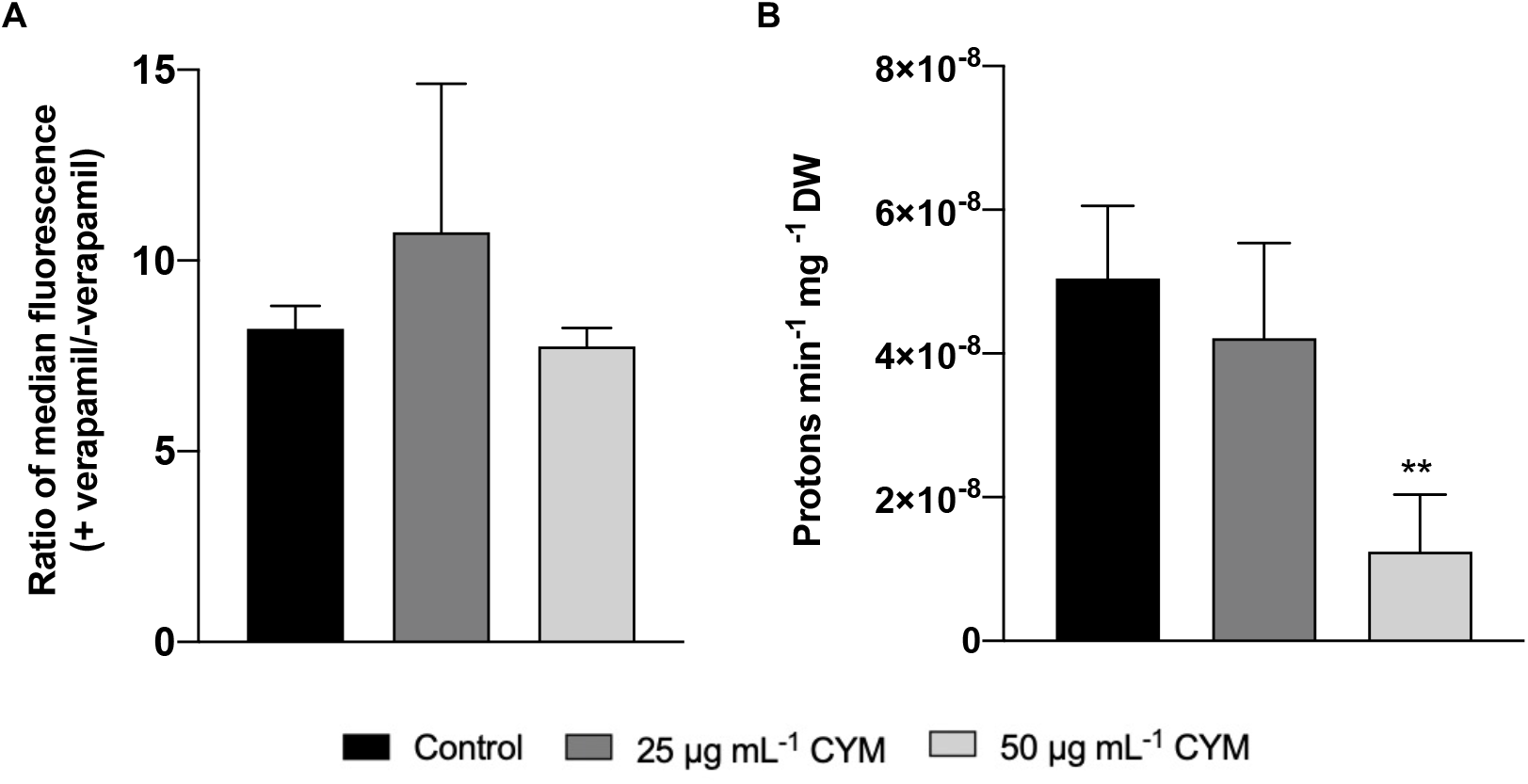
Effect of CYM on efflux pumps and Pma1p activity. BY4741 cells were treated for 8 h in the absence (control) or presence of 25 or 50 μg mL^-1^ CYM. **(A)** For evaluation of efflux pumps, cells were collected, treated with verapamil or solvent and stained with calcein AM. Data are presented as the ratio of median of the green fluorescence intensity between verapamil-treated and untreated cells (+ verapamil/ - verapamil). **(B)** Pma1p proton pumping activity was assessed by measuring H+ efflux triggered by glucose addition. Cells were collected and the pH adjusted to 5.5. Afterwards, 20 mM of glucose was added to induce proton extrusion by Pma1p. The proton extrusion activity was calculated from the maximum slope of the records and represents the rate of acidification. The data displayed are the mean ± standard deviation of three independent experiments. Asterisks (**, P ≤ 0.01) depict significant differences relative to the control.

We therefore next evaluated if CYM-induced alterations in lipid rafts could affect the activity of the proton pumping H+-ATPase Pma1p, one of the most abundant integral membrane proteins of the yeast plasma membrane and a resident of lipid rafts (40,41). We indeed found that proton pumping was affected by CYM, as exposure of cells to 50 μg mL^-1^ of CYM for 8 h led to a reduction in proton movement out of cells (Figure 6B).

As Pma1p pumps protons from the cytosol to the exterior of the cell coupled to ATP hydrolysis, Pma1p inhibition could be a result of decreased intracellular ATP levels. However, our data revealed that exposure to CYM is associated with increased intracellular accumulation of ATP (Figure 7A), instead consistent with less ATP consumption as a result of Pma1p inhibition (Figure 6A).

**Figure 7.**
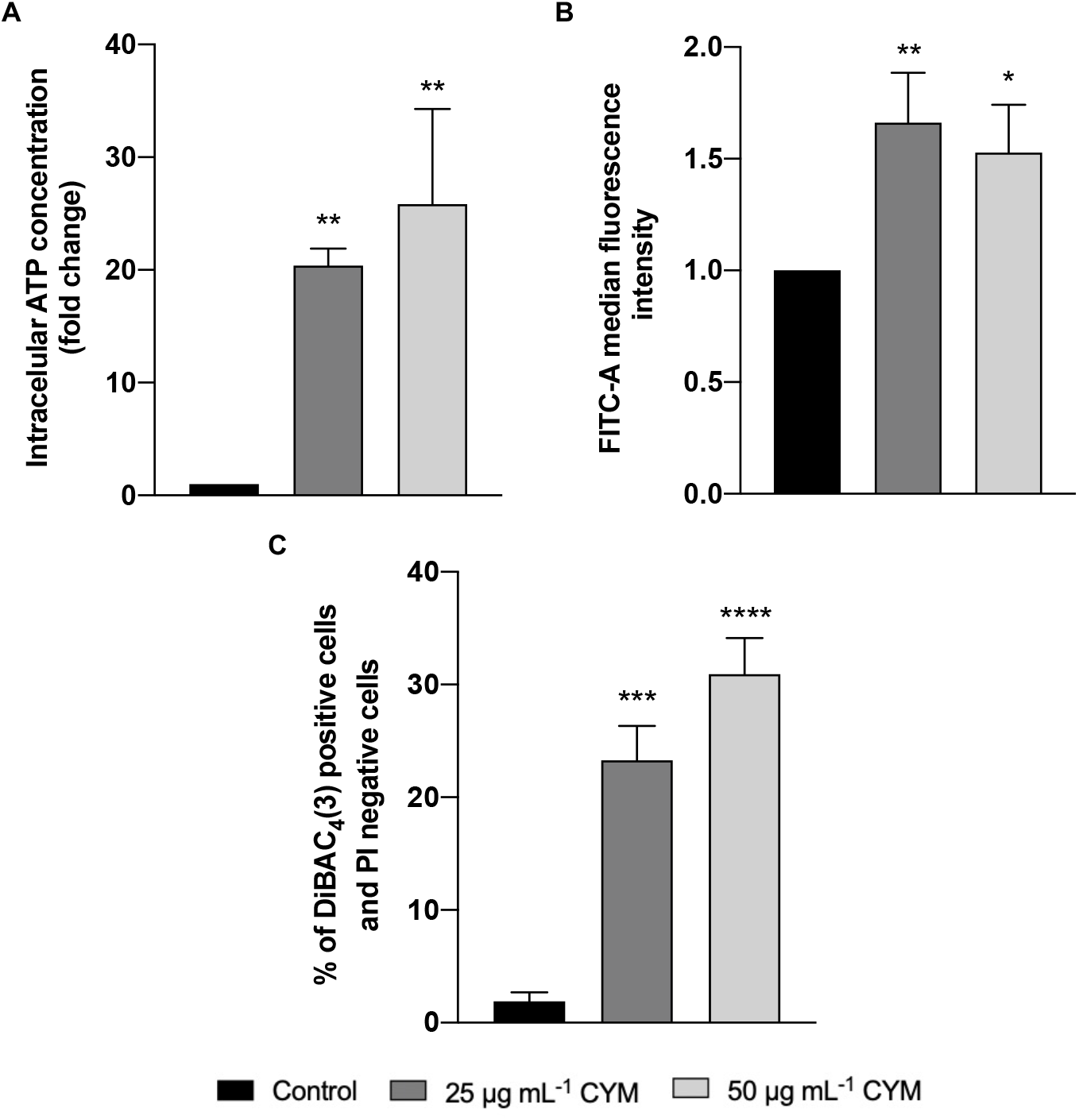
Effect of CYM on intracellular levels of ATP, vacuolar pH and membrane potential of BY4741 cells. Cells were treated for 8 h in the absence (control) or presence of 25 or 50 μg mL^-1^ of CYM. **(A)** Intracellular ATP levels were measured with the BacTiter-Glo™ Microbial Cell Viability kit. Values represent the increase in ATP levels expressed in fold change in comparison with the control. **(B)** Alterations in vacuolar pH were monitored using the CDCFDA probe. Data represent the green median fluorescence intensity normalized to control. **(C)** Plasma membrane potential and integrity were assessed through DiBAC_4_(3) and PI staining, respectively. Data represent the percentage of cells with green fluorescence (DiBAC_4_(3) positive cells) and without red fluorescence (PI negative cells). The data displayed are the mean ± standard deviation of three independent experiments. Asterisks (* ≤ 0.05; **, P ≤ 0.01; ***, P ≤ 0.001; ****, P ≤ 0.0001) depict significant differences relative to the control.

Pma1p activity, coupled with the activity of other pumps, such as the ATPase in the vacuole (V-ATPase), regulates intracellular pH homeostasis, critical for yeast cell survival. We therefore tested if CYM could affect the yeast vacuolar pH with the pH-sensitive probe CDCFDA. Figure 7B shows that CYM-treated cells exhibit a higher fluorescence signal than untreated cells (control), indicating that CYM also affects V-ATPase activity.

Since Pma1p proton pumping activity is also vital to generate an electrochemical gradient, we explored if CYM could cause alterations in plasma membrane potential using the fluorescent anionic probe DiBAC_4_(3). Simultaneously, and to distinguish between membrane depolarization and general membrane permeabilization, cells were stained with PI. We observed that the percentage of PI-negative/DiBAC_4_(3)-positive cells was higher in CYM-treated cells than in non-treated cells, indicating that CYM leads to plasma membrane depolarization prior to loss of plasma membrane integrity (Figure 7C).

## 4. Discussion

CYM is a synthetic fungicide used for foliar applications in a curative and protective manner, mainly against downy mildew diseases and late blight diseases (10,11). As it exhibits a relatively short period of activity when used alone due to its weak persistence (half-life of a few days, depending on the conditions) and mobile nature (adsorption coefficient, K_OC_ 39– 238) (42), it has been almost exclusively used with other fungicides, such as mancozeb and copper, to increase efficacy (12). Previous studies suggested that CYM can affect several biochemical processes such as growth, respiration, and synthesis of nucleic acids, but most of these alterations were considered secondary effects (13,14,43). However, despite widespread use, the CYM mode of action remains elusive, complicating efforts to assess how CYM affects different target and non-target organisms and highlighting the importance of careful consideration when using this fungicide. We therefore sought to use the yeast *S. cerevisiae* as model system for an in-depth investigation of the cellular and molecular mechanisms underlying CYM cytotoxicity.

We started by assessing sensitivity of yeast cells to CYM using a fermentable (glucose) or respiratory (galactose) substrate and observed that cells lost viability and plasma membrane integrity earlier in the former conditions. We therefore conclude that the effect of CYM on cell growth and viability is not dependent, at least not solely, on active respiration, and as such it must have additional effects. Therefore, further characterization of the effects of CYM in yeast cells were pursued using glucose as the carbon source.

To gain insight into the possible effects of CYM at the molecular level, we performed two different and complementary screens, transcriptomics and proteome localization. Among other hits, an effect on membrane transporter systems stood out in both screens. On one hand, there was up-regulation of genes encoding membrane transporters, and multiple membrane proteins appeared in an intracellular localization after CYM treatment. Moreover, several proteins appeared in the vacuole in response to CYM, which could be an indication of autophagy. A starvation phenotype with potential autophagy induction was also suggested by the categories enriched in transcriptomics analysis, such as a decrease in synthesis of most amino acids. Autophagy is considered a pro-survival mechanism that can be triggered by starvation and chemical stress (44), leading to vacuolar degradation of intracellular components (45,46). However, it also plays a critical role in normal development and cell homeostasis with intricate relations with cell metabolism, growth control, balance between cell survival and cell death, as well as ageing (47). We observed only a very tenuous band of free GFP in all conditions tested, which is consistent with basal levels of autophagy and thus disproving that CYM could induce autophagy.

As mentioned above, analysis of the enrichment in GO terms from both screens revealed that CYM affects genes associated with the plasma membrane. Consequently, and due to the importance of plasma membrane composition and organization to proper cellular function, we hypothesized that exposure to CYM could lead to plasma membrane alterations. We found that CYM changes plasma membrane organization by inducing intracellular accumulation of ergosterol and perturbing ergosterol-rich lipid rafts, plasma membrane nanodomains enriched in ergosterol and sphingolipids (48). We observed a similar pattern in membrane perturbation of yeast cells treated with the azole fungicide tebuconazole (49), known for inhibiting ergosterol biosynthesis. However, the alterations in lipid raft distribution induced by CYM were not accompanied by increased expression of *ERG* genes, in contrast to what was previously described for tebuconazole (49). Taken together, our results suggests that CYM disturbs ergosterol-rich lipid rafts without affecting ergosterol biosynthesis. Since deletion of *END3*, an essential gene for endocytosis, did not prevent intracellular accumulation of ergosterol, we hypothesize that CYM may impair the sorting of newly synthetized ergosterol to lipid rafts on the plasma membrane. However, it is also possible that CYM leads to increase endocytosis of plasma membrane proteins through a yet uncharacterized mechanism.

Alterations in lipid rafts can affect a myriad of cellular functions, eventually affecting cell fitness and viability, as these domains are associated with several biologically important proteins engaged in the regulation of ion homeostasis (Na^+^, K^+^), intracellular pH (H^+^), nutrient transport, mating processes, drug efflux and stress responses (40). We therefore first assessed if CYM could lead to a decrease in the activity of efflux pumps, which could result in increased intracellular accumulation of CYM and consequent loss of viability. It would also be consistent with a sensitization to other compounds, and would help explain the efficacy of CYM in mixed formulations. However, we did not find any differences on the activity of drug efflux pumps of CYM-treated cells. We next hypothesized that CYM may inhibit the activity of the plasma membrane proton ATPase Pma1p, one of the most abundant raft proteins. Pma1p generates and maintains an electrochemical gradient across the plasma membrane, which is vital for yeast cell survival since it is used for the uptake of inorganic ions and nutrients and to regulate the intracellular pH. Pma1p generates the electrochemical gradient by pumping protons from the cytosol to the exterior of the cell associated to ATP hydrolysis (50,51). Indeed, we found that CYM reduces the proton pumping activity of Pma1p. One option could be that CYM inhibits this pump, directly or indirectly through lipid raft alterations, and/or increased endocytosis, or that it indirectly leads to intracellular depletion of ATP, in turn inhibiting Pma1p activity. We found that CYM-treated cells have a higher intracellular accumulation of ATP, consistent with the first hypothesis. In addition to intracellular accumulation of ATP, which may not be deleterious *per se*, Pma1 inhibition can also lead to increased intracellular levels of H+, affecting plasma membrane potential and intracellular pH homeostasis. We found that CYM led to a decrease in the plasma membrane potential of yeast cells, again suggesting that CYM may function as a Pma1 inhibitor. We also observed that CYM leads to vacuolar alkalinization, suggesting that CYM also affects V-ATPase activity.

As an essential fungal protein, Pma1p has been a proposed target for the development of novel effective antifungal medications. One study previously screened a small molecule library for the ability of compounds to inhibit Pma1p, to identify compounds that could subsequently lead to plasma membrane depolarization and display antifungal activity (52). Other antifungal compounds with different properties have also been described to lead to a similar phenotype, such as lactoferrin and edelfosine, which also display anticancer activity. Both cause alterations in sterol organization at the plasma membrane, loss of Pma1p from lipid rafts and intracellular acidification in *S. cerevisiae* cells (28,53,54), and have been proposed as therapies to aid in the fight against drug-resistant fungal infections. It is conceivable that a similar mode of action for CYM could prove useful in its function as an agrochemical. CYM, which is cytotoxic, could first decrease the levels of target organisms, but plasma membrane alterations would also weaken surviving targets and render them more susceptible to other compounds. This could explain the use of CYM in joint formulations as an effective susceptibility enhancer of the effect of the substance that accompanies it, optimizing its short life.

## 5. Conclusion

Our work identifies for the first time the plasma membrane as one of the targets of CYM and proposes a mode of action underlying its antifungal activity: perturbation of lipid raft organization and inhibition of Pma1p and V-ATPase activity, leading to a decrease in plasma membrane potential and intracellular acidification, followed by cell death. These findings therefore contribute to a better understanding of the mechanism of action of CYM and its potential as a fungitoxic agent.

## Acknowledgements

We thank Yeynit Asraf and Zohar Gazi for work in high-content screening. Investigation by the authors has been supported by national funds (Portuguese Science Foundation, FCT) via the institutional programs supporting CBMA (UIDB/04050/2020, DOI: 10.54499/UIDB/04050/2020) and ARNET (LA/P/0069/2020), and funding to Susana Chaves DOI:10.54499/DL57/2016/CP1377/CT0026. Filipa Mendes was supported by a PhD scholarship from FCT (SFRH/BD/147574/2019). The robotic system of the Schuldiner laboratory was purchased through the kind support of the Blythe Brenden-Mann Foundation. Maya Schuldiner is an incumbent of Dr. Gilbert Omenn and Martha Darling Professorial Chair in Molecular Genetics.

## CRediT author statement

**Filipa Mendes:** Conceptualization, Data curation, Formal analysis, Investigation, Validation, Visualization, Writing - original draft, Writing – review & editing. **Hadar Meyer:** Data curation, Investigation, Supervision. **Leslie Amaral:** Formal analysis, Investigation. **Bruno B. Castro, Maria João Sousa, Maya Schuldiner and Susana R. Chaves:** Conceptualization, Funding acquisition, Project administration, Resources, Supervision, Validation, Writing - original draft, Writing - review & editing. All authors read and approved the manuscript.

## Footnotes

1 https://www.fao.org/faostat/en/#data/RP (assessed 18-10-2023)

